# Bridging micro and macro: accurate registration of the BigBrain dataset with the MNI PD25 and ICBM152 atlases

**DOI:** 10.1101/561118

**Authors:** Yiming Xiao, Jonathan C. Lau, Taylor Anderson, Jordan DeKraker, D. Louis Collins, Terry Peters, Ali R. Khan

## Abstract

Brain atlases that encompass detailed anatomical or physiological features are instrumental in the research and surgical planning of various neurological conditions. Magnetic resonance imaging (MRI) has played important roles in neuro-image analysis while histological data remain crucial as a gold standard to guide and validate such analyses. With cellular-scale resolution, the BigBrain atlas offers 3D histology of a complete human brain, and is highly valuable to the research and clinical community. To bridge the insights at macro- and micro-levels, accurate mapping of BigBrain and established MRI brain atlases is necessary, but the existing registration is unsatisfactory. The described dataset includes co-registration of the BigBrain atlas to the MNI PD25 atlas and the ICBM152 2009b atlases (symmetric and asymmetric versions) in addition to manual segmentation of the basal ganglia, red nucleus, and hippocampus for all mentioned atlases. The dataset intends to provide a bridge between insights from histological data and MRI studies in research and neurosurgical planning. The registered atlases, anatomical segmentations, and deformation matrices are available at: https://osf.io/xkqb3/.

## Background & Summary

Brain atlases are essential tools in neuroimage analysis and in neurosurgery, where they provide the reference to help navigate the anatomical and physiological features of the brain. While the foundational histology-derived atlases, such as Talairach^1^ and Schaltenbrand^2^ atlases established the seminal brain-based coordinate system for neurological navigation, their application was somewhat limited by the lack of accurate 3D reconstruction. The development of magnetic resonance imaging (MRI) has allowed sophisticated computational algorithms^3-5^ to reveal structural and functional variations in living brains due to neurological developments and disorders. Often averaged from multiple subjects, the newer MRI brain atlases^6-9^ provide high-quality anatomical and physiological information, which can be mapped to an individual’s brain to facilitate further analyses. Despite the advancements to improve resolution, MRI signals remain at macroscopic resolutions. The BigBrain atlas^10^ is a 3D digitized model of a human brain at a near-cellular 20 micrometer resolution. It is a unique tool to help integrate cytoarchitectural knowledge with MRI insights to study brain functions and to define anatomical structures that can be difficult to image in MRI (e.g., the subthalamic nucleus) for clinical practice and research. To bridge histological data with MRI, an accurate mapping between BigBrain and MRI brain atlases is necessary. Previously, a nonlinear registration between BigBrain and the ICBM152-2009b symmetric brain template^8^ was provided at <u>bigbrain.loris.ca</u>. This co-registration was achieved by deforming BigBrain with an inverted intensity profile to a population-averaged T1 map that is co-registered to the MNI space, with the SyN algorithm^11^ and cross-correlation similarity metric. However, this anatomical alignment, especially for subcortical structures, is not satisfactory, likely due to the discrepancy in tissue contrasts between BigBrain and the T1 map. As a result, more accurate alignment is greatly beneficial for various studies and surgical planning.

The ICBM152 brain atlas dataset^8^, from the Montreal Neurological Institute (MNI) is one of the most influential tools in neuroimage analysis. In total, MRI brain scans of 152 young adults at 1.5T were recruited to build the multi-contrast atlas, which includes T1w, T2w, and PDw contrasts, as well as probabilistic tissue maps and brain structural labels. After the initial edition with affine registration, the 2009 edition using group-wise nonlinear registration provides unbiased representation of the brain anatomy with sharp details. For this edition, both symmetric and asymmetric atlases were offered at the resolutions of 0.5×0.5×0.5mm^3^ (ICBM2009b) and 1×1×1mm^3^ (ICBM2009c).

As both natural ageing and neurological disorders can influence the anatomical features of the brain (e.g., tissue atrophy), population-specific atlases^7-9^ are created to ensure the quality of neuroimage analysis and surgical planning. Aiming to facilitate the research and surgical treatment of Parkinson’s disease (PD), the MNI PD25 population-averaged atlases^7,12^ were constructed from 3T MRI scans^13^ of 25 PD patients, and contain five different image contrasts, including T1w, T2*w, T1–T2* fusion, phase, and an R2* map. The special T1-T2* fusion atlas has the general T1w contrast for most of the brain while preserving the subcortical structures, such as the basal ganglia, red nucleus, and dentate nucleus as shown in typical T2*w contrast, which often suffers from susceptibility artefacts near the cortical surface. Furthermore, the dataset is co-registered with a digitized histological atlas with 123 structures^14^ and probabilistic tissue maps.

In the dataset described here, we introduce an accurate nonlinear registration of BigBrain to the symmetric and asymmetric versions of ICBM2009b atlas and the MNI PD25 atlas. As suggested by earlier studies^15,16^, T1w-to-T1w registration can be sub-optimal for subcortical structures (e.g. subthalamic nucleus) that are nearly invisible in T1w MRI, we employed a two-stage multi-contrast registration procedure with the PD25 space as the medium, as shown in *Fig.1*. The proposed method takes advantage of the similar contrast between BigBrain and PD25 T1-T2* fusion atlases, and a synthetic T2w PD25 template to ensure the structural alignment between MRI atlases since T2*w and T2w MRIs differ greatly in the contrast of neuroanatomical structures, with T2*w particularly sensitive to iron in tissues. For the atlases involved (BigBrain and all MRI atlases), the basal ganglia, red nucleus, thalamus, amygdala, and hippocampus were manually segmented at high resolution as additional shape priors to ensure atlas-to-atlas warping^17^ and to help validate the final registration outcomes. We expect the described dataset to greatly benefit the clinical and research community.

**Figure 1.**
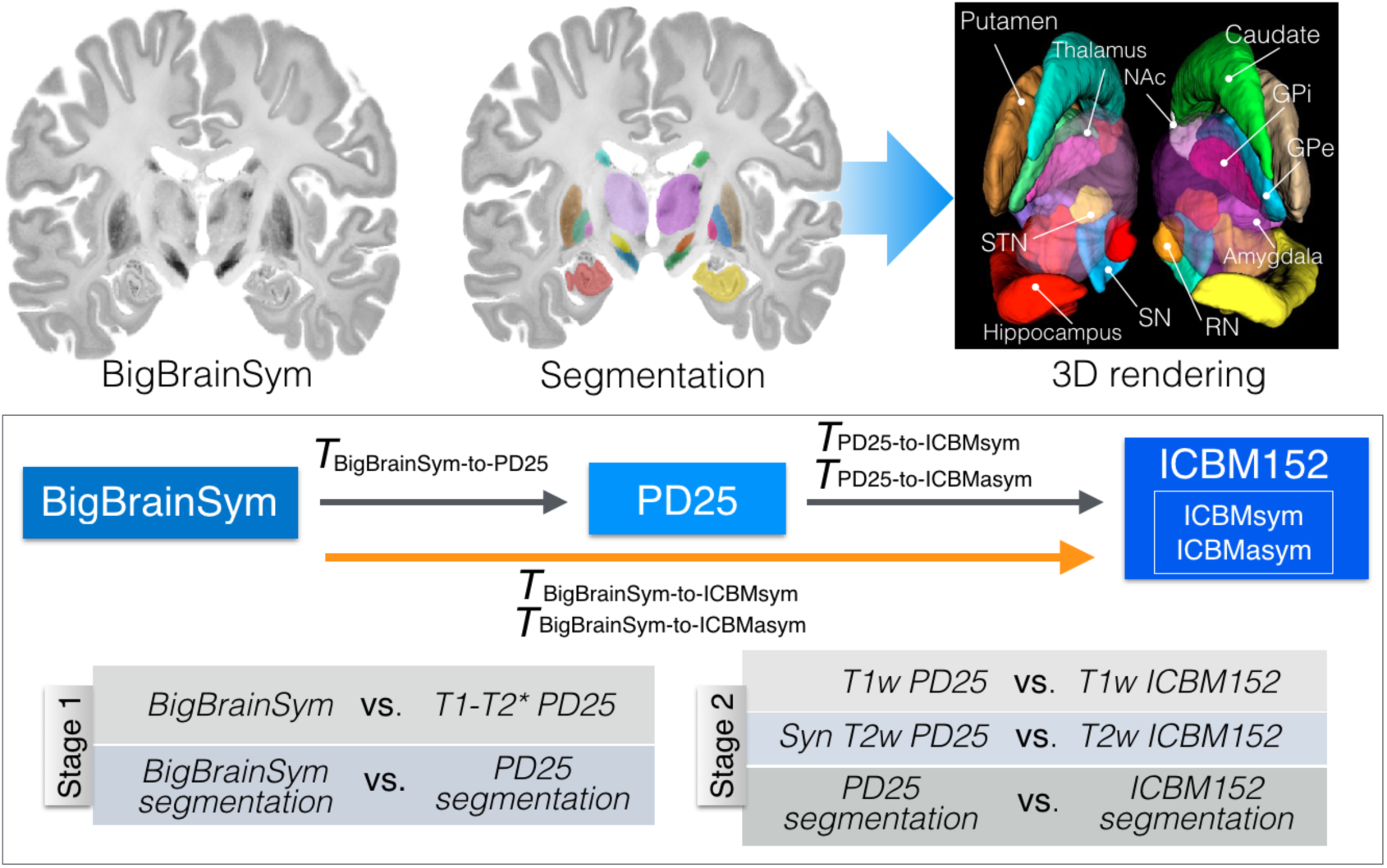
Manual segmentation of the subcortical structures for the BigBrainSym atlas by the author YX with hippocampus shown as semi-transparent (top row) and the schematic of the two-stage registration strategy for BigBrain-to-ICBM152 alignment (bottom row). For the registration strategy, the contrast pairs used at each registration stage are also listed.

## Methods

### Manual segmentation

Manual segmentations were used to facilitate atlas-to-atlas registration and validate the registration results. This approach ensures the optimal structural overlap in multi-modal registration^17^, and thus reduces the potential loss that is propagated to atlas-to-subject mapping. To simplify the notations for all atlases involved in this article, we refer to the symmetric and asymmetric versions of ICBM152 2009b release as *ICBMsym* and *ICBMasym*, respectively, and use the name *BigBrainSym* to call the original co-registered BigBrain atlas to the ICBM152 space as provided in the BigBrain 2015 release. To aid the readers, the list of short names for different atlases is provided in Table 1. Here, eleven pairs of subcortical structures were manually segmented at 0.3×0.3×0.3mm^3^ resolution for *BigBrainSym*, and at 0.5×0.5×0.5mm^3^ for the MNI PD25 and the ICBM2009b symmetric and asymmetric atlases. These structures include the putamen, caudate nucleus, globus pallidus pars externa (GPe), globus pallidus pars interna (GPi), nucleus accumbens (NAc), amygdala, thalamus, red nucleus (RN), substantia nigra (SN), subthalamic nucleus (STN), and hippocampus. The segmentation was performed using ITK-SNAP (itksnap.org) with the left and right side labelled separately. While the RN and the basal ganglia structures were labelled by the author TA and revised by YX, who is experienced in brain anatomy, the rest were completed by YX. Here, the hippocampus segmentation follows the protocol employed by DeKraker et al.^18^, and the amygdala and NAc labels follow the protocol by Pauli et al.^6^. The full list of segmented structures and their associated label numbers are provided in Table 2. To inspect the quality of manual segmentation, the same set of anatomical structures were also manually segmented by a co-author (JD) with expertise in neuroanatomy and physiology, for BigBrainSym, PD25, and ICBMsym at 0.5×0.5×0.5mm^3^ resolution. The segmentations were compared against those by YX using Dice coefficient, and the results are shown in Table 3. For BigBrainSym, the segmentation by YX was downsampled to 0.5×0.5×0.5mm^3^ with nearest-neighbourhood interpolation for comparison.

**Table 1.** Descriptions for all the abbreviations of atlases employed.

|  | Description |
| --- | --- |
| **BigBrainSym** | The registration of BigBrain to ICBM152 symmetric space provided as in the 2015 BigBrain data release |
| **PD25** | The MNI PD25 atlas for a Parkinson's disease cohort |
| **ICBMsym** | The symmetric version of ICBM152 2009b atlas |
| **ICBMasym** | The asymmetric version of ICBM152 2009b atlas |

**Table 2.** Label numbers with the corresponding nuclei for subcortical segmentation of all atlases.

| Label number | Nucleus | Label number | Nucleus |
| --- | --- | --- | --- |
| 1 | Left red nucleus | 2 | Right red nucleus |
| 3 | Left substantia nigra | 4 | Right substantia nigra |
| 5 | Left subthalamic nucleus | 6 | Right subthalamic nucleus |
| 7 | Left caudate | 8 | Right caudate |
| 9 | Left putamen | 10 | Right putamen |
| 11 | Left globus pallidus externa | 12 | Right globus pallidus externa |
| 13 | Left globus pallidus interna | 14 | Right globus pallidus interna |
| 15 | Left thalamus | 16 | Right thalamus |
| 17 | Left hippocampus | 18 | Right hippocampus |
| 19 | Left nucleus accumbens | 20 | Right nucleus accumbens |
| 21 | Left amygdala | 22 | Right amygdala |

**Table 3.** Inter-rater structural segmentation measured as Dice coefficient.

|  | BigBrainSym | ICBMsym | PD25 |
| --- | --- | --- | --- |
| L Red Nucleus | 0.94 | 0.96 | 0.93 |
| R Red Nucleus | 0.95 | 0.96 | 0.91 |
| L SN | 0.92 | 0.91 | 0.94 |
| R SN | 0.94 | 0.91 | 0.93 |
| L STN | 0.87 | 0.89 | 0.92 |
| R STN | 0.90 | 0.89 | 0.91 |
| L Caudate | 0.94 | 0.97 | 0.92 |
| R Caudate | 0.94 | 0.97 | 0.94 |
| L Putamen | 0.97 | 0.96 | 0.97 |
| R Putamen | 0.97 | 0.96 | 0.96 |
| L GPe | 0.96 | 0.96 | 0.98 |
| R GPe | 0.95 | 0.96 | 0.98 |
| L GPi | 0.93 | 0.96 | 0.92 |
| R GPi | 0.93 | 0.94 | 0.93 |
| L Thalamus | 0.98 | 0.98 | 0.98 |
| R Thalamus | 0.98 | 0.98 | 0.98 |
| L hippocampus | 0.94 | 0.97 | 0.96 |
| R hippocampus | 0.94 | 0.97 | 0.96 |
| L Nucleus Accumbens | 0.96 | 0.97 | 0.93 |
| R Nucleus Accumbens | 0.96 | 0.97 | 0.95 |
| L Amygdala | 0.96 | 0.96 | 0.97 |
| R Amygdala | 0.95 | 0.96 | 0.97 |

Here, the Dice coefficient (κ) for assessing inter-rater variability is computed by:

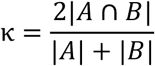

where A and B are two different segmentations, respectively, and | · | represents the number of voxels within the segmentation. A value of κ = 1 represents a prefect overlap, and no overlap gives a value of 0.

### Synthetic T2w PD25 template

As the MNI PD25 dataset primarily leverages the T2*w contrast to visualize the subcortical structures (e.g., the STN, RN and SN), direct mapping between the PD25 T1-T2* atlas and T2w MRI scans can be challenging due to differences in image contrasts. To facilitate the inter-contrast registration, a synthetic T2w PD25 atlas, *I*_*syn*−*T*2*w*_ was constructed as:

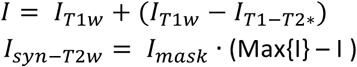

where *I*_*T*1*w*_ and *I*_*T*1−*T*2*_ are the T1w and T1-T2* PD25 atlases, *I*_*mask*_ is the brain mask, and *Max{I}* is the maximum value within the image *I*. The resulting synthetic T2w PD25 atlas is shown in *Fig.2*, alongside the co-registered BigBrain atlas.

**Figure 2.**
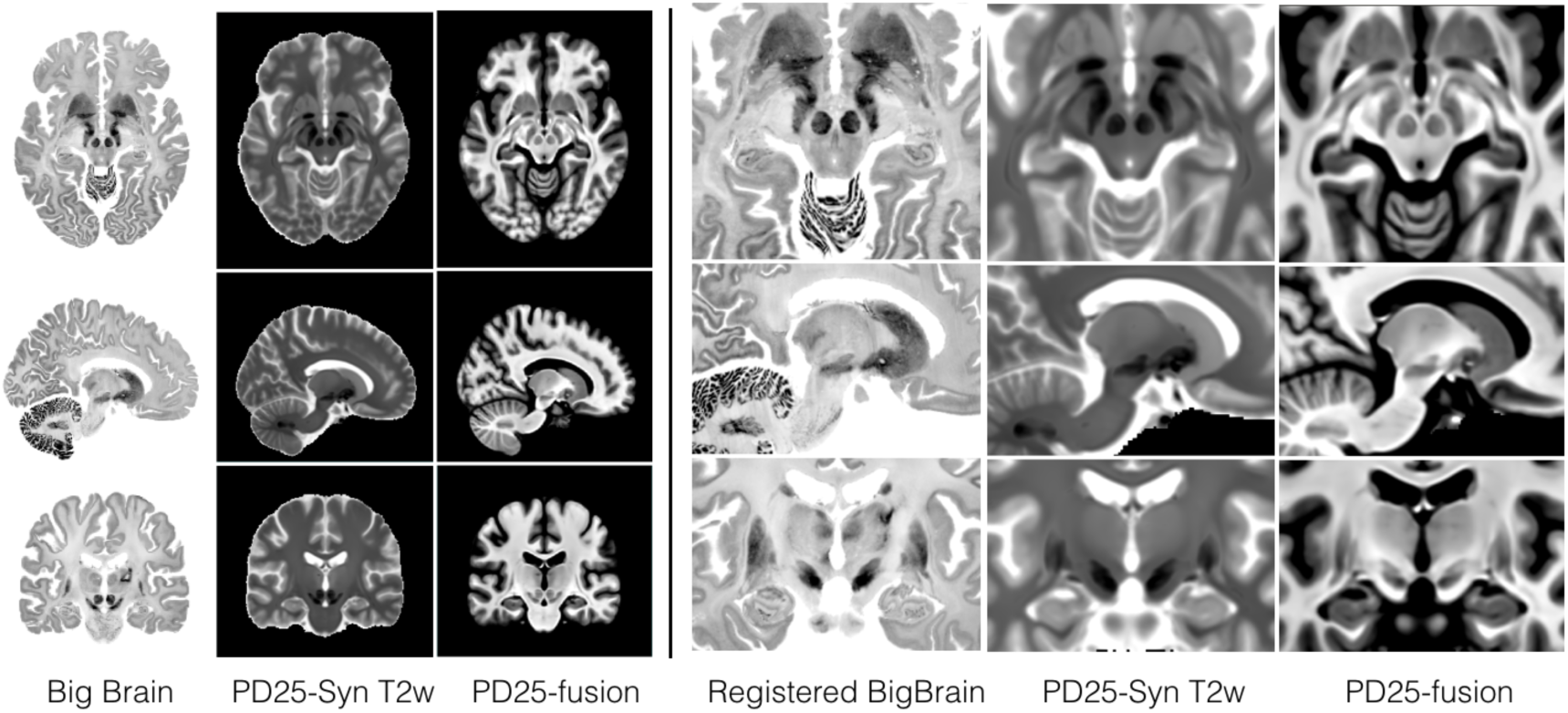
Comparison of the BigBrain atlas registered to PD25 atlas, T1-T2* fusion PD25 atlas, and synthetic T2w PD25 atlas with corresponding slices across images. The results are shown for the entire brain (left) and the deep brain region (right).

### Atlas registration

We employed *BigBrainSym* to initiate the atlas-to-atlas registration as it provided a good starting point. There are two main difficulties in mapping *BigBrainSym* to the ICBM152 atlases. First, the reconstructed histological volume has a unique and different appearance from the ICBM152 atlases, making accurate nonlinear registration with conventional image similarity metrics (e.g., mutual information) challenging. Second, tissue tear and distortion from histology handling created unrealistic morphology (e.g., excessive distortion in hippocampus) and artifacts (e.g., tear in right thalamus) that can adversely influence the mapping. To mitigate these issues, we implemented a two-stage multi-contrast strategy to warp between *BigBrainSym* and the ICBM152 atlases, by using the MNI PD25 space as an intermediate template and adding anatomical segmentations as shape priors to further guide the registration. More specifically, *BigBrainSym* was first nonlinearly registered to the PD25 space, which was then deformed to the *ICBMsym* or *ICBMasym* atlas. Lastly, the deformation fields from the two stages were concatenated, and used to resample the *BigBrainSym* to the ICBM152 space. For both stages, we used *antsRegistration* from the Advanced Normalization Tools (ANTs, stnava.github.io/ANTs) to achieve the image registration, and all images were processed in MINC2 format.

Taking advantage of the contrast similarity between *BigBrainSym* and the T1-T2* PD25 atlas, we used this pair of images to achieve the registration in the first stage. Inherited from the data in the native histological space, *BigBrainSym* contains a few problematic examples of anatomical morphology and artefacts. Besides those mentioned earlier, *BigBrainSym* also has an oversized pineal gland and tectum - likely from tissue stretching during histological processing. To cope with these, the pineal gland was removed from BigBrainSym for registration to avoid over-stretching of local deformation, which can adversely affect the overall registration, and the tectum was segmented in addition to the 11 subcortical structures in both atlases to constrain the registration. The multi-class segmentations were placed in one image for each atlas and blurred by a Gaussian kernel with a full-width-at-half-maximum (FWHM) of 0.5 mm. Finally, they were used jointly with the atlases during registration. Here, we used Mattes mutual information and cross-correlation for the atlas pair (weight=1) and the segmentations (weight=0.8), respectively.

As shown in Fig. 1, to map the PD25 space to the ICBM152 space, the T1w and T2w contrasts (synthetic T2w contrast for PD25), together with the subcortical segmentations, were jointly employed. Similar to the first stage, the labels of 11 subcortical structures were blurred by a Gaussian kernel with a FWHM=0.5mm. For each contrast pair, a cross-correlation metric was used, and weights of 1, 1, and 0.8 were assigned to T1w contrast, T2w contrast, and subcortical segmentations, respectively during registration cost function optimization.

In the two-stage registration strategy, PD25 was used as an intermediate volume due to a few considerations. First, compared with the previous approach that uses intensity inversion and a T1 map as the intermediate volume, the T1-T2* fusion contrast of PD25 has much closer resemblance to BigBrain, particularly for the subcortical structures. This can facilitate automatic registration. Second, BigBrain was derived from a 65-year-old healthy male, whose age is within the range of PD25 cohort (age=58±7 years). The difference in brain atrophy patterns in normal aging and Parkinson’s disease without cognitive impairment (target group for PD25) is relatively small^19^, and pair-wise nonlinear registration should sufficiently account for the anatomical differences. Lastly, we hope the resulting dataset can benefit both healthy and pathological population, as well as neurosurgical applications, such as deep brain stimulation (DBS). The inclusion of PD25 atlas, therefore, will be very beneficial.

### Atlas registration evaluation

The quality of atlas registration was assessed with two widely employed approaches: 1) anatomical landmark (fiducials) registration errors and 2) atlas-based subcortical segmentation accuracy. While the first metric evaluates the matching of distinct anatomical features, the latter validates the correspondence of subcortical structures. Both metrics were computed for *BigBrain-to-PD25, PD25-to-ICBM152* (symmetric and asymmetric versions), and finally *BigBrain-to-ICBM152* (symmetric and asymmetric versions) registration. Additionally, as a reference, we also calculated the two metrics between *BigBrainSym* and *ICBMsym*.

To assess the atlas alignment with landmark registration errors, we used the anatomical fiducials (AFIDs) framework introduced by Lau *et al*.^20^ (2019). For the framework, anatomical landmarks were selected by eight experienced raters for *BigBrainSym, ICBMsym*, and *ICBMasym*, and the final landmark coordinates at each location was obtained by averaging the results from all raters after filtering out outlier points. Following the same protocol, the final anatomical landmarks for the MNI PD25 atlas were produced by five experience raters based on the T1w atlas. The full details of the landmark picking protocols and the associated software can be found in the original AFIDs article^20^. Since for *BigBrainSym*, excessive tissue tear and distortion exists, we excluded the pineal gland and culmen from the original AFIDs protocol^20^ for registration validation. The 30 anatomical landmarks employed for registration validation are listed in Table 4, where the Euclidean distance between the transformed point and the target point was computed for each landmark location.

**Table 4.** Landmark registration errors (mm) for all registrations, as well as for the BigBrain vs. *ICBMsym* registration in BigBrain 2015 data release.

|  | BigBrainSym-to-PD25 | PD25-to-ICBMasym | PD25-to-ICBMsym | BigBrainSym-to-ICBMasym | BigBrainSym-to-ICBMsym | BigBrainSym<br>vs.ICBMsym |
| --- | --- | --- | --- | --- | --- | --- |
| AC | 0.73 | 0.26 | 0.19 | 0.72 | 0.62 | 0.74 |
| PC | 0.23 | 0.23 | 0.21 | 0.36 | 0.40 | 0.45 |
| infracollicular<br>sulcus | 1.62 | 1.46 | 1.52 | 0.74 | 1.39 | 6.36 |
| PMJ | 2.39 | 0.53 | 0.18 | 1.81 | 2.07 | 0.61 |
| superior<br>interpeduncular<br>fossa | 2.31 | 0.72 | 0.42 | 1.52 | 2.17 | 1.62 |
| R superior LMS | 2.00 | 0.36 | 0.37 | 1.61 | 2.00 | 1.23 |
| L superior LMS | 1.90 | 0.39 | 0.49 | 1.97 | 1.32 | 1.34 |
| R inferior LMS | 0.55 | 0.26 | 2.09 | 0.57 | 2.28 | 1.60 |
| L inferior LMS | 1.49 | 0.22 | 1.82 | 1.48 | 1.16 | 1.89 |
| intermamillary<br>sulcus | 0.50 | 0.29 | 0.52 | 0.75 | 0.61 | 0.52 |
| R MB | 0.63 | 0.30 | 0.51 | 0.66 | 0.30 | 1.13 |
| L MB | 0.43 | 0.58 | 0.92 | 0.79 | 0.80 | 1.15 |
| R LV at AC | 3.14 | 0.86 | 0.50 | 2.21 | 2.68 | 1.98 |
| L LV at AC | 4.61 | 2.09 | 1.77 | 2.47 | 2.85 | 2.05 |
| R LV at PC | 2.63 | 2.25 | 2.22 | 0.49 | 0.79 | 1.27 |
| L LV at PC | 2.07 | 2.84 | 1.49 | 2.12 | 1.20 | 1.31 |
| genu of CC | 0.64 | 0.29 | 0.68 | 0.81 | 0.83 | 0.65 |
| splenium of CC | 2.20 | 0.56 | 0.53 | 1.85 | 1.98 | 2.23 |
| R AL temporal<br>horn | 1.48 | 1.59 | 1.78 | 0.16 | 0.45 | 0.70 |
| L AL temporal<br>horn | 1.76 | 1.31 | 2.12 | 1.86 | 2.78 | 4.69 |
| R superior AM<br>temporal horn | 3.86 | 1.19 | 0.54 | 2.86 | 3.68 | 0.89 |
| L superior AM<br>temporal horn | 4.89 | 1.89 | 1.63 | 3.01 | 3.32 | 1.68 |
| R inferior AM<br>temporal horn | 3.43 | 2.24 | 2.34 | 1.75 | 1.53 | 0.83 |
| L inferior AM<br>temporal horn | 4.13 | 1.53 | 2.07 | 2.35 | 1.88 | 1.88 |
| R indusium<br>griseum origin | 2.24 | 1.26 | 1.95 | 1.30 | 0.58 | 1.21 |
| L indusium<br>griseum origin | 2.76 | 0.31 | 2.48 | 2.67 | 1.39 | 0.74 |
| R ventral<br>occipital horn | 5.46 | 1.22 | 1.23 | 4.58 | 4.57 | 2.54 |
| L ventral<br>occipital horn | 7.00 | 1.58 | 1.38 | 5.09 | 5.29 | 5.88 |
| R olfactory<br>sulcal fundus | 0.81 | 0.55 | 0.55 | 0.72 | 0.66 | 2.62 |
| L olfactory<br>sulcal fundus | 1.42 | 0.84 | 0.45 | 2.16 | 1.60 | 3.06 |
| Mean±sd | 2.31±1.66 | 1.00±0.74 | 1.17±0.76 | 1.71±1.16 | 1.77±1.25 | 1.83±1.47 |

The Dice coefficient (κ) metric was used to evaluate the quality of volumetric overlap between the native manual segmentation and the corresponding labels warped from another atlas. For smaller structures, such as the midbrain nuclei, values greater than 0.7 are usually accepted as good segmentations while for larger structures, values above 0.8 are preferred.

## Data Records

The complete dataset includes deformed atlases, subcortical segmentations, and inter-atlas spatial transformations. More specifically, we supply the BigBrain atlas deformed to the PD25, *ICBMsym*, and *ICBMasym* atlases at three different resolutions (0.3×0.3×0.3mm^3^, 0.5×0.5×0.5mm^3^, and 1×1×1mm^3^). The subcortical segmentations (see Table 2) were included for BigBrainSym at 0.3×0.3×0.3mm^3^, and for PD25, ICBMsym, and ICBMasym at 0.5×0.5×0.5mm^3^. All these image volumes are made available in both MINC2 and NIfTI-1 formats. The script *mnc2nii* from MINC Toolkit (http://bic-mni.github.io) was used for image format conversion. Lastly, the dataset provides the nonlinear transformations for *BigBrainSym-to-PD25, PD25-to-ICBMsym, PD25-to-ICBMasym, BigBrainSym-to-ICBMsym*, and *BigBrainSym-to-ICBMasym* registrations. All these transformations are provided in MINC transformation format. The full dataset can be accessible at the Open Science Framework (OSF)^21^ (http://doi.org/10.17605/OSF.IO/XKQB3), as well as the main project page for the MNI PD25 atlases at <u>nist.mni.mcgill.ca/?p=1209</u>. In addition, all manually segmented label maps and brain atlases are also made available in BIDS format at OpenNeuro.org^22^ (https://openneuro.org/datasets/ds002016/versions/1.0.0).

## Technical Validation

### Anatomical landmark registration

Landmark registration errors were computed for all individual anatomical landmarks and for the five sets of atlas-to-atlas registrations involved in this dataset. The results are shown in Table 4. The calculated mean registration errors are 2.31, 1.00, 1.17, 1.71, and 1.77 mm for *BigBrainSym-to-PD25, PD25-to-ICBMasym, PD25-to-ICBMsym, BigBrainSym-to-ICBMasym*, and *BigBrainSym-to-ICBMsym*, respectively. For comparison, in Table 4, the results were also listed for BigBrain’s original registration to the symmetric ICBM152 space (*BigBrainSym* vs. *ICBMsym*). In general, the introduced two-stage registration strategy resulted in a slightly better mean registration error (1.77±1.25 mm) than the previous registration for *BigBrain* vs. *ICBMsym* (1.83±1.47 mm), but by performing a pair-wise Wilcoxon signed rank test, this difference is not significant (p=0.959). As mentioned in the original AFIDs article^20^, ventricular features (e.g., ventral occipital horns) generally have higher landmark placement errors, and landmark placement is also more difficult for individual subjects than population-averaged atlases. This can be even more challenging for histological data, where unnatural deformation and unique individual anatomical features exist. As shown in Table 4, the registration error varies for different anatomical landmarks, potential users should sufficiently consider this factor during their application of this co-registration.

When looking closer at all registration results, the averaged errors from MRI-to-MRI registrations were lower than those from BigBrain-to-MRI registrations. This is expected, since individual anatomical variability is more pronounced than group-averaged anatomy and inter-modality registration is more challenging. Also, in terms of landmark registration errors, *BigBrainSym* is better aligned with the ICBM152 space than PD25 for certain landmarks, likely due to better population representativeness with a larger cohort and the fact that the BigBrain landmarks were tagged within the *BigBrainSym* atlas^20^, potentially making the these landmark placement more biased towards the ICBM152 space.

### Subcortical structural segmentation

Dice coefficients were calculated for 11 pairs of anatomical structures as listed in Table 5, and the results are listed for all atlas-to-atlas alignments. The mean Dice coefficients were computed at 0.94, 0.94, 0.94, 0.93, 0.93 for *BigBrainSym-to-PD25, PD25-to-ICBMasym, PD25- to-ICBMsym, BigBrainSym-to-ICBMasym*, and *BigBrainSym-to-ICBMsym*, respectively. In contrast to a mean κ value of 0.77 from the original warping of BigBrain to the symmetric ICBM152 space, the new strategy greatly improved the subcortical alignment for all structures of interest. By adding manual labels in multi-contrast registration, the alignment of subcortical anatomy was relatively consistent across different registrations. This helps ensure the quality of atlas-to-subject registration for future investigations by reducing the accuracy loss in multi-modal atlas-to-atlas warping, and the same approach^17^ was employed earlier for histology-to-MRI registration. Although the manual segmentations were also used for validation, the improved deformation is substantial in terms of Dice coefficient measurements for the subcortical structures, and it is also evident by visual inspection in *Fig.3*, particularly for the regions annotated with colored arrows.

**Figure 3.**
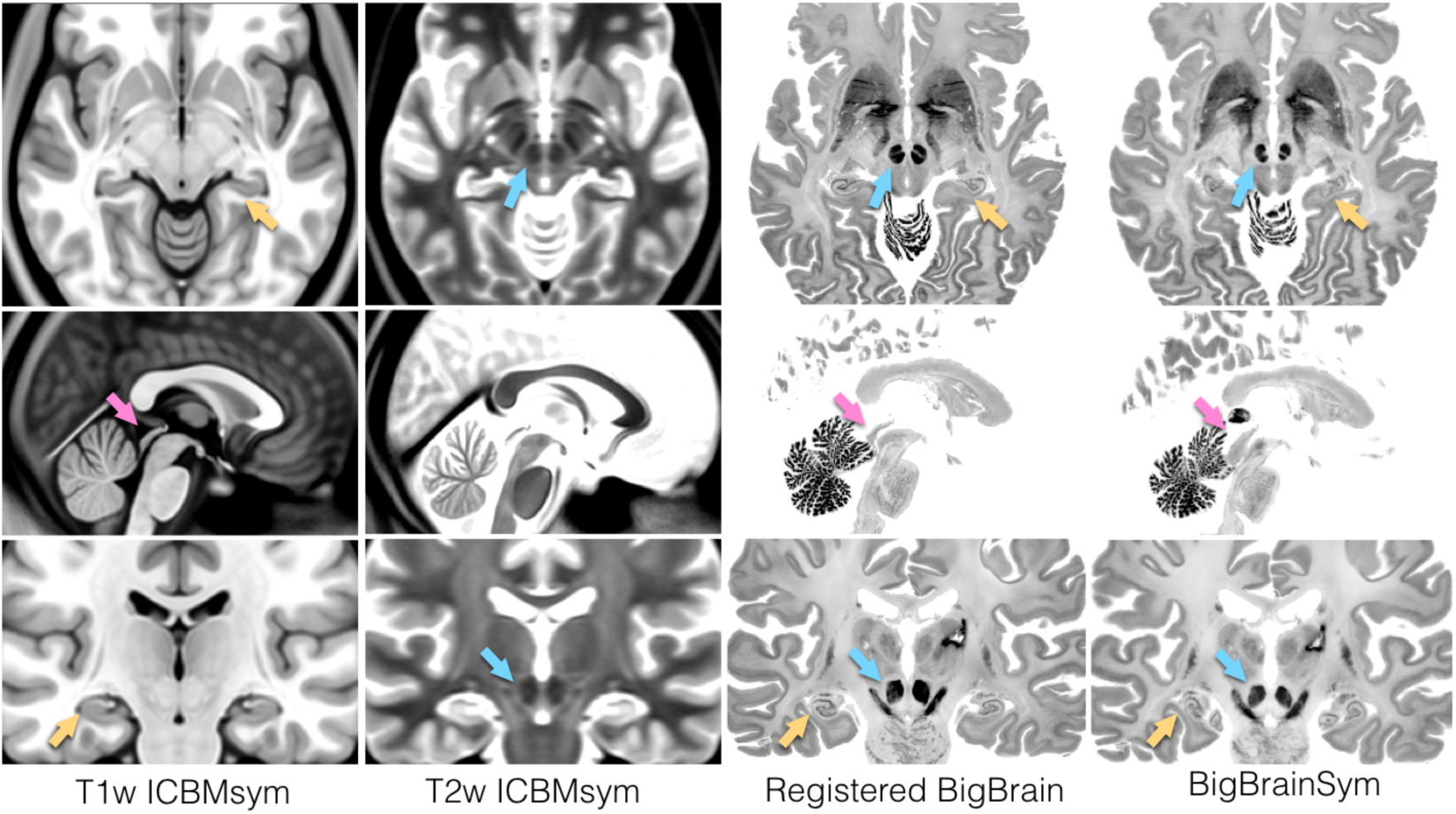
Comparison of *ICBMsym*, new co-registered BigBrain to *ICBMsym*, and *BigBrainSym*. Each row corresponds to the same slice within each atlas, and each column shows the axial, sagittal, and coronal views of an atlas. The visible improvements in red nucleus, tectum, and hippocampus are annotated with blue, pink, and yellow arrows in different anatomical views, respectively. Note that for in the new registration of BigBrain, the pineal gland was removed.

**Table 5.** Dice coefficients of subcortical structures for all atlas-to-atlas registrations, as well as for the BigBrain vs. *ICBMsym* registration in BigBrain 2015 data release. In the table, L = left and R = right.

|  | BigBrainSym-to-PD25 | PD25-to-ICBMasym | PD25-to-ICBMsym | BigBrainSym-to-ICBMasym | BigBrainSym-to-ICBMsym | BigBrainSym<br>vs.ICBMsym |
| --- | --- | --- | --- | --- | --- | --- |
| L Red Nucleus | 0.91 | 0.90 | 0.91 | 0.89 | 0.89 | 0.76 |
| R Red Nucleus | 0.93 | 0.92 | 0.91 | 0.90 | 0.90 | 0.70 |
| L SN | 0.94 | 0.91 | 0.91 | 0.91 | 0.92 | 0.79 |
| R SN | 0.92 | 0.92 | 0.91 | 0.92 | 0.92 | 0.77 |
| L STN | 0.91 | 0.85 | 0.84 | 0.87 | 0.86 | 0.76 |
| R STN | 0.88 | 0.87 | 0.86 | 0.88 | 0.88 | 0.73 |
| L Caudate | 0.97 | 0.97 | 0.97 | 0.97 | 0.97 | 0.91 |
| R Caudate | 0.97 | 0.97 | 0.97 | 0.97 | 0.96 | 0.88 |
| L Putamen | 0.97 | 0.97 | 0.97 | 0.97 | 0.97 | 0.91 |
| R Putamen | 0.97 | 0.97 | 0.97 | 0.96 | 0.97 | 0.86 |
| L GPe | 0.93 | 0.92 | 0.93 | 0.92 | 0.92 | 0.74 |
| R GPe | 0.90 | 0.93 | 0.93 | 0.89 | 0.90 | 0.71 |
| L GPi | 0.90 | 0.92 | 0.92 | 0.90 | 0.90 | 0.72 |
| R GPi | 0.93 | 0.92 | 0.92 | 0.92 | 0.93 | 0.73 |
| L Thalamus | 0.98 | 0.98 | 0.98 | 0.98 | 0.98 | 0.88 |
| R Thalamus | 0.98 | 0.98 | 0.98 | 0.98 | 0.98 | 0.89 |
| L hippocampus | 0.94 | 0.95 | 0.95 | 0.93 | 0.93 | 0.62 |
| R hippocampus | 0.94 | 0.95 | 0.95 | 0.94 | 0.94 | 0.75 |
| L Nucleus Accumbens | 0.94 | 0.95 | 0.95 | 0.94 | 0.94 | 0.55 |
| R Nucleus Accumbens | 0.94 | 0.94 | 0.94 | 0.94 | 0.94 | 0.60 |
| L Amygdala | 0.95 | 0.96 | 0.96 | 0.95 | 0.94 | 0.81 |
| R Amygdala | 0.95 | 0.96 | 0.96 | 0.96 | 0.95 | 0.84 |

## Usage Notes

We provide the refined deformation matrices in MINC transformation format to comply with the existing releases of the BigBrain dataset. The linear and original nonlinear transformations between the BigBrain atlas in native histological space and the symmetric ICBM152 space are available at ftp://bigbrain.loris.ca/BigBrainRelease.2015/3D_Volumes/MNI-ICBM152_Space.

## Acknowledgements

The author Y. Xiao is supported by BrainsCAN and CIHR postdoctoral fellowships for this work.

## Author contributions

Y.X. was responsible for the study concept, image registration, subcortical segmentation of ICBM152 and PD25 atlases, inspection of segmentations, image processing, registration validation, and writing of the manuscript. J.C.L. organized the study to curate the anatomical landmarks for all brain atlases, generated the final fiducial landmarks for registration validation, and contributed to the writing of the manuscript. T.A. performed manual segmentation of basal ganglia and red nucleus for the BigBrainSym atlas. J. D. inspected the hippocampus segmentations for all atlases and provided manual segmentation for inter-rater variability assessment. D.L.C., T.P., and A.R.K supervised the study, provided materials and funding, and contributed to the editing of the manuscript.

## References

1 Talairch, J. T. P. (Thieme Medical Publishers, New York, 1988).

2 Schaltendbrand G.; Wahren, W. Atlas for stereotaxy of the human brain. 2nd edn, (Year Book Medical Publishers, 1977).

3 Whitwell, J. L. Voxel-based morphometry: an automated technique for assessing structural changes in the brain. J Neurosci 29, 9661–9664 (2009).

4 Khan, A. R. et al. Biomarkers of Parkinson’s disease: Striatal sub-regional structural morphometry and diffusion MRI. Neuroimage Clin (2018).

5 Vemuri, P., Jones, D. T. & Jack, C. R., Jr. Resting state functional MRI in Alzheimer’s Disease. Alzheimers Res Ther 4, 2 (2012).

6 Pauli, W. M., Nili, A. N. & Tyszka, J. M. A high-resolution probabilistic in vivo atlas of human subcortical brain nuclei. Sci Data 5, 180063 (2018).

7 Xiao, Y. et al. Multi-contrast unbiased MRI atlas of a Parkinson’s disease population. Int J Comput Assist Radiol Surg 10, 329–341 (2015).

8 Fonov, V. et al. Unbiased average age-appropriate atlases for pediatric studies. Neuroimage 54, 313–327 (2011).

9 Liang, P. et al. Construction of brain atlases based on a multi-center MRI dataset of 2020 Chinese adults. Sci Rep 5, 18216 (2015).

10 Amunts, K. et al. BigBrain: an ultrahigh-resolution 3D human brain model. Science 340, 1472–1475 (2013).

11 Avants, B. B., Epstein, C. L., Grossman, M. & Gee, J. C. Symmetric diffeomorphic image registration with cross-correlation: evaluating automated labeling of elderly and neurodegenerative brain. Med Image Anal 12, 26–41 (2008).

12 Xiao, Y. et al. A dataset of multi-contrast population-averaged brain MRI atlases of a Parkinsons disease cohort. Data Brief 12, 370–379 (2017).

13 Xiao, Y., Beriault, S., Pike, G. B. & Collins, D. L. Multicontrast multiecho FLASH MRI for targeting the subthalamic nucleus. Magn Reson Imaging 30, 627–640 (2012).

14 Chakravarty, M. M., Bertrand, G., Hodge, C. P., Sadikot, A. F. & Collins, D. L. The creation of a brain atlas for image guided neurosurgery using serial histological data. Neuroimage 30, 359–376 (2006).

15 Xiao, Y. et al. in Proceedings of the Third international conference on Information Processing in Computer-Assisted Interventions 135–145 (Springer-Verlag, Pisa, Italy, 2012).

16 Xiao, Y. et al. Investigation of morphometric variability of subthalamic nucleus, red nucleus, and substantia nigra in advanced Parkinson’s disease patients using automatic segmentation and PCA-based analysis. Hum Brain Mapp 35, 4330–4344 (2014).

17 Ewert, S. et al. Toward defining deep brain stimulation targets in MNI space: A subcortical atlas based on multimodal MRI, histology and structural connectivity. Neuroimage 170, 271–282 (2018).

18 DeKraker, J., Ferko, K. M., Lau, J. C., Kohler, S. & Khan, A. R. Unfolding the hippocampus: An intrinsic coordinate system for subfield segmentations and quantitative mapping. Neuroimage 167, 408–418 (2018).

19 Weintraub, D. et al. Neurodegeneration across stages of cognitive decline in Parkinson disease. Arch Neurol 68, 1562–1568 (2011).

20 Lau, J. C. et al. A framework for evaluating correspondence between brain images using anatomical fiducials. Hum Brain Mapp (2019).

21 Xiao, Y. et al. Accurate registration of the BigBrain dataset with the MNI PD25 and ICBM152 atlases. doi:10.17605/OSF.IO/XKQB3 (2019).

22 Xiao, Y. et al. BigBrainMRICoreg. OpenNeuro, doi:10.18112/openneuro.ds002016.v1.0.0 (2019).

